# BISoN: A Bayesian Framework for Inference of Social Networks

**DOI:** 10.1101/2021.12.20.473541

**Authors:** Jordan D. A. Hart, Michael N. Weiss, Daniel W. Franks, Lauren J. N. Brent

**Affiliations:** Centre for Research in Animal Behaviour, University of Exeter, UK; Center for Whale Research, Friday Harbour, WA, USA; Departments of Biology and Computer Science, University of York, UK

## Abstract

1. Social networks are often constructed from point estimates of edge weights. In many contexts, edge weights are inferred from observational data, and the uncertainty around estimates can be affected by various factors. Though this has been acknowledged in previous work, methods that explicitly quantify uncertainty in edge weights have not yet been widely adopted, and remain undeveloped for many common types of data. Furthermore, existing methods are unable to cope with some of the complexities often found in observational data, and do not propagate uncertainty in edge weights to subsequent statistical analyses.
2. We introduce a unified Bayesian framework for modelling social networks based on observational data. This framework, which we call *BISoN*, can accommodate many common types of observational social data, can capture confounds and model effects at the level of observations, and is fully compatible with popular methods used in social network analysis.
3. We show how the framework can be applied to common types of data and how various types of downstream statistical analyses can be performed, including non-random association tests and regressions on network properties.
4. Our framework opens up the opportunity to test new types of hypotheses, make full use of observational datasets, and increase the reliability of scientific inferences. We have made example R scripts available to enable adoption of the framework.

## Introduction

Social network analysis is arguably one of the most popular frameworks in the study of sociality (Croft et al., 2016). In many scientific contexts, network connections (edges) are inferred from observational data, from human observers recording social interactions between monkeys, to biologgers capturing proximity events between people. The type of data used to infer networks and the manner in which the data are collected affects both the interpretation of social network analyses and their accuracy (Whitehead, 2008).

Networks are usually constructed by taking a normalised measure of sociality, such as the proportion of sampling periods each pair spends engaged in a social behaviour (e.g. the Simple Ratio Index; SRI). These normalised measures ensure that pairs that are observed for longer are not erroneously determined to be more social (Whitehead, 2008). This is important because observing social events can be challenging and uniform sampling over all pairs is not always possible (Croft et al., 2010). Though normalised measures of sociality will not be biased by sampling time, they will be accompanied by varying levels of certainty. For example, edge weights will be treated as a certain 0.5 for both a case where individuals have been seen together once and apart once, and equally where individuals have been together 100 times and apart 100 times. Despite there being considerably more certainty around the values of the latter example than the former, methods to estimate and propagate uncertainty through social network analyses remain largely underdeveloped.

In this paper we introduce BISoN: a general unified Bayesian framework for modelling social network data. BISoN captures uncertainty in edge weights, making full use of available data; propagates uncertainty through downstream analyses; and can control for social and non-social effects at any level (individual, dyad, group, observation, etc). Any type of social network analysis can be conducted within our framework, including dyadic and nodal regressions, non-random edge weight tests, and estimation of structural network properties. The BISoN framework comprises a) an *edge weight model*, dependent on data type, that builds edge weights and networks with uncertainty from empirical data, and b) downstream analyses that use estimated edge weights to take into account uncertainty in the network. We will focus on how our framework can be applied to animal systems, but the underlying principle can be applied to any type of network analysis where network edges are inferred. The BISoN framework presents an opportunity to generate reliable, flexible, and rich scientific inference in the study of social systems.

## BISoN - Bayesian Inference of Social Networks

In the following section we outline the three edge weight models we have developed for binary, count, and duration data (see Table 1 for definitions of these types of data). For brevity the models are presented without detailing any specific priors, but when fitting these models, priors should always be specified, and the choice of prior should depend on the context and the question in hand (van de Schoot et al., 2021). These edge weight models can be used to generate estimates for network edge weights *ω_ij_*. We describe how to obtain edge weight estimates for binary, count and duration data below. Once estimates for edge weights *ω_ij_* are obtained, they can be used in downstream analyses. We will briefly outline three types of analyses that propagate uncertainty through the analysis. These analyses are: 1) testing for non-random edge weights, 2) dyadic regression, and 3) nodal regression. When presenting the downstream analyses, for brevity we will refer to them in the context of the count data model, but these downstream analyses are freely applicable to all types of data. The models we present here are simple examples that can be extended and refined in many ways. The Bayesian nature of this framework means that these methods will be useful and reliable, providing they are used appropriately in the context of the data, the scientific question, and with appropriate diagnostic tools (Kruschke, 2015; McElreath, 2020). We have provided example code at: https://www.github.com/JHart96/BISON_examples.

**Table 1:** Definitions of the three types of social observational data we have developed models for. Examples of standard edge weight interpretations are included in the third column.

| Data type | Description | Edge interpretation |
| --- | --- | --- |
| Binary | The presence or absence of a social event for each dyad is recorded in each sampling period. | Probability of (or proportion of time) engaging in a social event in a fixed period of time. |
| Count | The number of social events for each dyad is recorded in each sampling period. | Rate of occurrence of social events per unit time. |
| Duration | The amount of time each dyad spends engaged in a social event is recorded in each sampling period. | Proportion of time spent engaging in a social event. |

### Quantifying uncertainty in edge weights

#### Edge weight model: Binary data

To demonstrate the notion behind BISoN, we will first describe the edge weight model for the case of binary data, where presence or absence of a social event per fixed sampling period is recorded. First, note that for binary data, edge weights are commonly defined as:

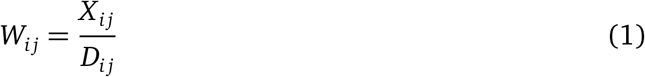

where *X_ij_* is the number of social events that occurred between *i* and *j*, and *D_ij_*. is the number of sampling periods where it is considered possible for a social event to have been observed between *i* and *j*. The exact definition will depend on context, but this may often be the number of periods in which at least one of the individuals of the dyad was seen.

The edge weight is therefore equivalent to the probability of *i* and *j* engaging in a social event in any given sampling period. This can also be interpreted as an estimate of the proportion of time *i* and *j* spend engaging in a social event. Let us refer to this probability (or proportion) as *p_ij_*. Without making any additional assumptions beyond that of the standard SRI calculation, an equivalent formulation is:

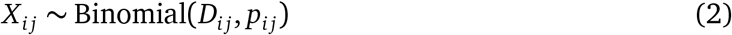

Note that the maximum likelihood estimate for *p_ij_* in this process is *X_ij_/D_ij_* = *W_ij_*. The binomial process is the natural process for multiple Bernoulli trials, which makes this formulation a natural extension of point estimates for edge weights. In a Bayesian context, the parameter *p_ij_* is treated as a random variable, and inherently captures any uncertainty around its value. A similar formulation was used by Farine and Strandburg-Peshkin (2015) to estimate uncertainty over edge weights using a beta conjugate prior over the *p_ij_* parameters (as detailed in Fink (1997)).

Social data can sometimes be influenced by non-social effects at the observation level, for example varying levels of habitat visibility depending on location, variance in observer reliability, or time of day can all affect the recorded observations of social events. These effects cannot be modelled by aggregating social events *X_ij_* at the dyad level. To solve this, we propose to model these effects by decomposing the binomial process over the aggregated observations to distinct Bernoulli processes over unaggregated observations, as follows:

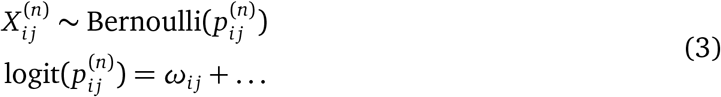

where 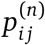 is the probability of a social event occurring between *i* and *j* in the *n*-th observation of *i* and *j*, and *ω_ij_* is the edge weight between *i* and *j*. The ellipsis represents additional terms that can be included (if desired) to model various effects. The term 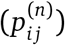 can be seen as analogous to the predictor terms in regression models. In the location-dependent visibility example, including an observation-level location effect will account for this visibility effect. When additional effects are included, the edge weight will become an estimate of relative sociality in the presence of non-social effects. We use a linear combination of effects in these examples for simplicity, but the effects could equally be modelled as non-linear functions if deemed necessary in context. Alternatively dyad- or individual-level effects can also be included to separate out different types of social effect.

The binary data model can also be used for group-based (gambit) data (Franks et al., 2010). If being used for group data, an additional term in the model can be used to account for nonindependence within group observations. See the supplementary material for further discussions on these points.

#### Edge weight model: Count data

The same concept as above can also be applied to the case of count data, where counts of social events between individuals per sampling period is recorded. First, note that for count data, point estimates of edge weights are often defined in the same way as Equation 1, but where *X_ij_* is the total number of social events that occurred between *i* and *j*, and *D_ij_* is the total amount of time a social event could have been observed between *i* and *j*.

The edge weight is therefore equivalent to the rate at which *i* and *j* engaged in a social event per unit time. Let us refer to this rate as *λ_ij_*. When events occur with a fixed rate, the number of events expected per unit time is described by the Poisson process (Wasserman, 2004). Since we assume that the rate of events characterises edge weights well, then without making any additional assumptions, we can model social events as:

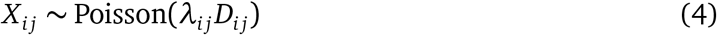

where *λ_ij_* is the underlying rate parameter. When modelled in a Bayesian framework, this provides a natural way to model the uncertainty around edge weights because the maximum likelihood estimate for *λ_ij_* is *X_ij_/D_ij_* = *W_ij_* (Held and Sabanés Bové, 2014).

Again, this model doesn’t allow observation-level effects to be modelled, so we must decompose the aggregated data into the original sampling periods. Fortunately, the sum of Poisson-distributed random variables is also described by a Poisson distribution. This means that the full model can take a similar form to the previous model:

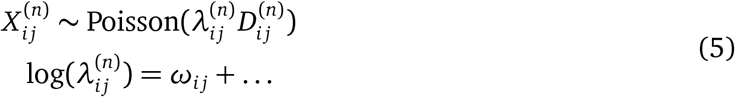

where 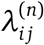 is the rate at which social events occur between *i* and *j* in the *n*-th observation of *i* and *j*, 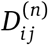 is the length of time of the *n*-th sampling period for the dyad, and *ω_ij_* is again the edge weight between *i* and *j*. As above, the ellipsis represents potential additional effects that could be included in the model if necessary. This is discussed in a later section and in the supplementary material.

#### Edge weight model: Duration data

The edge weight model for duration data is slightly more complex than those of the binary and count data models, because it models two different types of data simultaneously: the duration of social events, and the frequency with which they occur. With duration data, the edge weight is again calculated using Equation 1, but where *X_ij_* is the amount of time *i* and *j* were seen engaged in a social event, and *D_ij_* is the maximum amount of time *i* and *j* could have been seen engaged in a social event. The total amount of time *X_ij_* is the sum of times from *K_ij_* social events. The edge weight is then modelled as the proportion of time *t_ij_* each dyad could engage in a social event that is actually spent engaging in the social event:

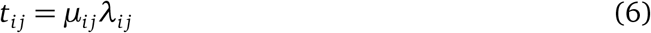

where *μ_ij_* is the mean event duration for the dyad and *λ_ij_* is the mean rate of events per unit time for the dyad. Inter-arrival times in a Poisson process are distributed according to the exponential distribution, so we use this to model the duration of social events once they have started. The count of events can again be considered to be a Poisson process. Bringing these two assumptions together gives the following model (see supplementary material for the full derivation):

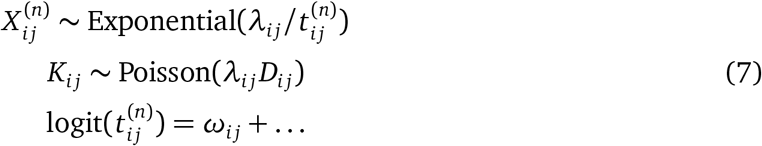

The mean event time *μ_ij_* is modelled inherently in 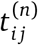 and can be recovered from the fitted model afterwards using *μ_ij_* = *ω_ij_/λ_ij_*.

### Accounting for social and non-social effects

By modelling social events at the observation-level rather than the dyad-level, it is possible to account for observation-level effects that may influence the estimated edge weights. An example of this is where the study population might move through various locations with different visibilities, leading to location-dependent missed social events. Assuming that variation in sociality between location isn’t of direct interest to the researcher, we can include a location effect in the model. An example of how this can be achieved in the count model is:

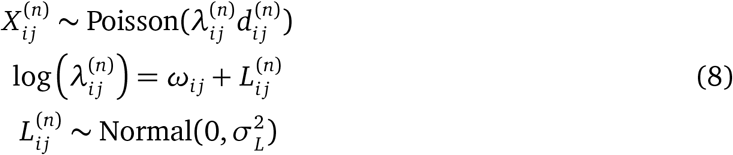

where 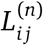 is a location effect corresponding to the location of the social events observed for the dyad *ij* in the *n*-th observation period. The location effect has an adaptive normal prior centered at zero (often known as a random effect). The location effect is designed to capture the differences in visibility at different locations. In theory, the effect of visibility on edge weights will always be negative, but we choose to model it withouts bounds. This is because the true visibilities in different locations are unknown, so the edge weights cannot be estimated exactly, only proportionally. It is therefore unnecessary to place an upper bound on the visibility effect, making the model easier to fit.

Depending on the assumed causal process that generates observations, it won’t always be possible to control for non-social effects in this way, as the social effect of interest may partially be borne out through apparent non-social effects, such as space use. For example indviduals may genuinely be more social in some locations than others, and controlling for location would remove the social effect of interest. A discussion on the the considerations around modelling non-social effects is included in the supplementary material.

### Downstream analysis with uncertainty

The previous section has shown how uncertainty can be estimated over edge weights. Once these estimates have been obtained, it is then possible to conduct various types of statistical analysis on the network while propagating uncertainty through the entire analysis. In this section we will discuss how several common types of social network analysis can be applied to social networks constructed with BISoN edge weight models. In particular we outline how three specific types of network analysis can be conducted: 1) testing for non-random edge weights, 2) dyadic regression, and 3) nodal regression.

#### Testing for non-random edge weights

BISoN enables a Bayesian test analogous to the classic non-random association test developed by Bejder et al. (1998), and originally proposed as the ‘chance sociogram’ by Moreno and Jennings (1938). At its core, this test asks whether the structure of an observed network is an artefact of sampling or if there is genuine variation in edge weights between individuals. If the structure of an observed network is simply due to sampling, then there is no variation in edge weight, and each dyad is equally likely to engage in social events as all other dyads. In BISoN, this is equivalent to

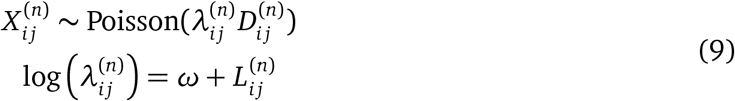

where *ω* is now a single edge weight parameter shared between all dyads, indicating no variation in underlying edge weights between dyads. This model constitutes the null hypothesis *H*_0_ of random edge weights. This can be compared to the alternative hypothesis *H*_1_ of nonrandom edge weights, where the *ω* parameter is replaced with a dyad-level parameter *ω_ij_*, allowing each dyad to exhibit a different edge weights. In a Bayesian context, these two models (*H*_0_: *ω_ij_* = ω vs *H*_1_: *ω_ij_* ≠ *ω*) can be compared using the *Bayes factor*. Instead of attempting to reject the null model, as in null hypothesis significance testing, the Bayes factor quantifies the support for or against the hypotheses. The Bayes factor in favour of *H*_1_ over *H*_0_ is defined as:

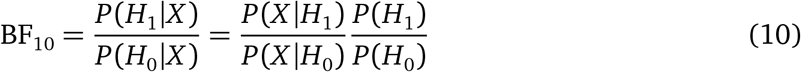

where *P*(*H*_1_|*X*) and *P*(*H*_0_|*X*) denote the probability of the alternative hypothesis and the null hypothesis given the data *X* respectively. *P*(*X*|*H*_1_) and *P*(*X*|*H*_0_) are known as the marginal likelihoods, and can be estimated numerically from a fitted model (Gronau et al., 2017). The Bayes factor can be directly interpreted as the relative likelihood of the competing hypotheses. For example, a Bayes factor of 5 in favour of *H*_1_ over *H*_0_ would indicate that *H*_1_ is 5 times more likely than *H*_0_ given the data and model. A useful property of Bayes factors is that they can quantify support for or against hypotheses, and can even be used to compare multiple hypotheses.

The original test from Bejder et al. (1998) is extendable to test for non-random associations within categories (such as sex combination or age difference). However, it requires continuous variables such as age difference to be discretised into categories. Discretising variables is generally not ideal, and removes information, weakens statistical power, and can lose useful insights (Altman and Royston, 2006). Our proposed test can account for variables such as age without discretisation. This can be achieved by adding variables to the predictor in the following way:

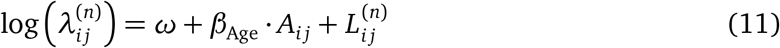

where *β*_Age_ is a parameter describing the effects of age on the log rate of social events, and *A_ij_* is the age difference for the dyad *i j*. In addition to modelling continuous variables, this approach to the non-random edge weight testing also makes it possible to compare multiple competing hypotheses of edge structure, model the effects of different dyad-level covariates, and compare hypotheses relating to global network metrics such as network density or clustering coefficients.

### Dyadic regression

A common type of analysis in studies of social networks is to consider factors that affect edge weights. This type of analysis is often called dyadic regression, where explanatory factors such as sex or age difference are regressed against edge weight. The edge weights in dyadic regression are non-independent, as variables such as age difference or sex difference are inherently linked to individual-level attributes. This means that effects due to age or sex in individual *i* affect all dyads that connect *i*. This non-independence can be controlled for by including node-level effects in the regression (Tranmer et al., 2014).

Using the count data example discussed earlier, we propose to conduct dyadic regression using a standard regression model of the form:

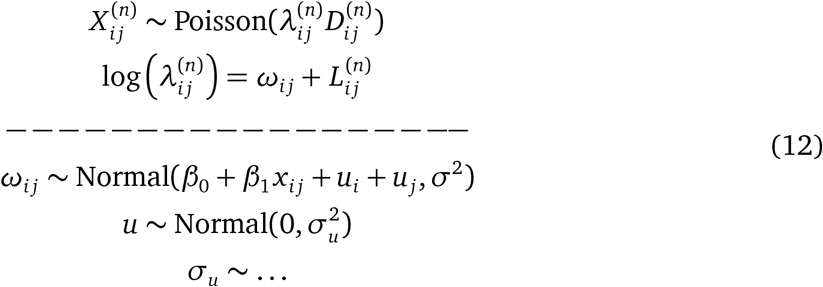

where *β*_0_ is the intercept parameter, *β*_1_ is the slope parameter, *u*_i_ are parameters accounting for the effect of node membership on edge weight, *σ* is the standard deviation of the residuals, and *σ_u_* is a hyperprior for the standard deviation of the node membership effect (undefined here to avoid suggesting a default prior). The *u* parameters are used to capture non-independence of dyads, and are sometimes known as multi-membership effects (Tranmer et al., 2014). The dashed line indicates that the model can be fit in two separate parts: the first part being the edge weight model, and the second part being the dyadic regression model. It is not strictly necessary to separate out the model like this, but doing so makes it possible to fit the edge weight model once and conduct multiple types of analysis on it afterwards without fitting the entire model again. The *β*_0_ and *β*_1_ parameters can be interpreted in the same way as usual regression coefficients, and will be accompanied with posterior distributions describing plausible values of the coefficients.

The model shown here is intended as a minimal example of a simple linear regression with node-level effects to account for the non-independence of edges. The regression can be extended to support different family distributions, hierarchical effects, non-linearities, and more. We have included examples of diagnostics and some extensions in the example code.

### Nodal regression

The final common type of network analysis we will cover here is nodal regression, where a regression is performed to analyse the relationship between a nodal network metric (such as centrality) and nodal traits (such as age and sex). These analyses are usually used to assess how network position depends on various biological factors, but can also be used where network position is a predictor. Since node metrics are derivative measures of the network, uncertainty in edge weights should ideally propagate through to node metrics, and on to coefficients in regression analyses, giving an accurate estimate of the total uncertainty in inferred parameters. The core of the nodal regression is similar to dyadic regression, taking the form:

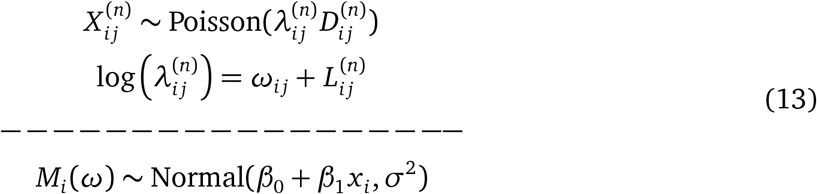

where *β*_0_ is the intercept parameter, *β*_1_ is the slope parameter, and *σ* is the standard deviation of the residuals. *M_i_*(*ω*) denotes the node metric estimates when applied to the edge weights estimated in the top part of the model.

Calculating node metrics within the model may present a practical challenge when using standard model fitting software, as many node metrics can not be described purely in terms of closed-form equations (Freeman, 1978). In this case, splitting up the model along the dashed line becomes an important step, as it allows the edge weights to be estimated using a standard piece of software, and leaves the regression part of the model to be fit using a sampler that supports numeric estimates of network centralities. As part of the code we have made available, we have written a custom Metropolis-Hastings sampler that maintains the joint distribution of edge weights, rather than sampling edge weights independently. This ensures that the effect of any structure in the observation data (such as location effects) is maintained and propagated through to the node metric estimates, and subsequently to regression coefficient estimates.

## Example: Synthetic dataset

To demonstrate our framework, we present an example based on a synthetic dataset of social events simulating count data. The synthetic dataset was generated by assigning underlying undirected edge weights to 28 dyads corresponding to all possible pairs of 8 individuals living in a closed social group. Individuals 1 - 4 were assigned to a treatment condition and individuals 4 - 8 were assigned to a control. Dyads between two individuals within the treatment condition were assigned a higher edge weight than other dyads, to induce an effect of treatment condition on centrality. Social events were simulated from a Poisson point process based on both the underlying edge weight and an observation-level, non-social effect of location. There were 6 possible locations for observations of social events that were used to simulate the effect of varying visibilities in different locations. Between 10 and 50 observation periods of fixed length (arbitrary time units) were simulated for each dyad. Full details of the simulation can be found in the example code.

The synthetic dataset describes the count of events for each observation period, therefore the appropriate model for this data is the count model. The full model was defined as:

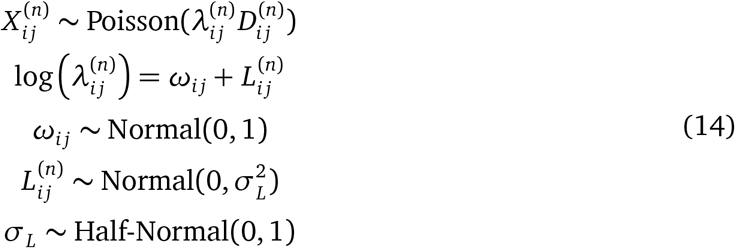

where 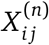 is the count of social events between *i* and *j* in their *n*-th observation period, 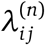 is the social event rate from the corresponding period, 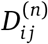 is the length of the corresponding sampling period (fixed to 1 here for simplicity), *ω_ij_* is the dyad-level edge weight, and 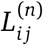 is an observation-level effect corresponding to one of the 6 locations. The prior distributions were chosen using a prior predictive check to ensure the parameters only take on biologically plausible values. The model was implemented in Stan, and fitted in R, using the package *Rstan* (R Core Team, 2020; Stan Development Team, 2020).

The model fit was checked visually by examining the chains and computing the Gelman-Rubin 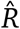 convergence statistic (Gelman and Rubin, 1992). Once the model fit was verified, the parameter estimates of edge weights *ω_ij_* between dyads were extracted. Figure 2A shows a visualisation of the network based on edge weight estimates *ω_ij_*, where the bands around edges are proportional to the widths of the 95% credible intervals. At this point the estimated network can be treated much the same as a standard network of point estimates, as long as care is taken to preserve uncertainty through subsequent analyses. To demonstrate the power of BISoN for conducting downstream analyses, we conducted a regression analysis where the response was eigenvector centrality and the predictor was a categorical variable with two categories, representing the treatment condition. To improve model fit, the response was standardised by subtracting the mean and dividing by the standard deviation. The model specification for the regression analysis was:

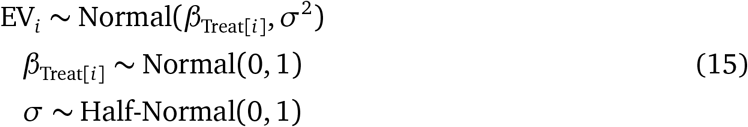

where EV*_i_* denotes the standardised eigenvector centrality for the *i*-th node, *β*_Treat_[*_i_*] is the effect parameter corresponding to the treatment condition (control or treatment) for the *i*-th sample, and *σ* is the standard deviation of the residuals. The regression model was fit using our custom Metropolis-Hastings sampler and chain convergence was checked visually and with the 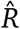 statistic. Posterior predictive checks were used to ensure the model fitted the data well.

**Figure 1:**
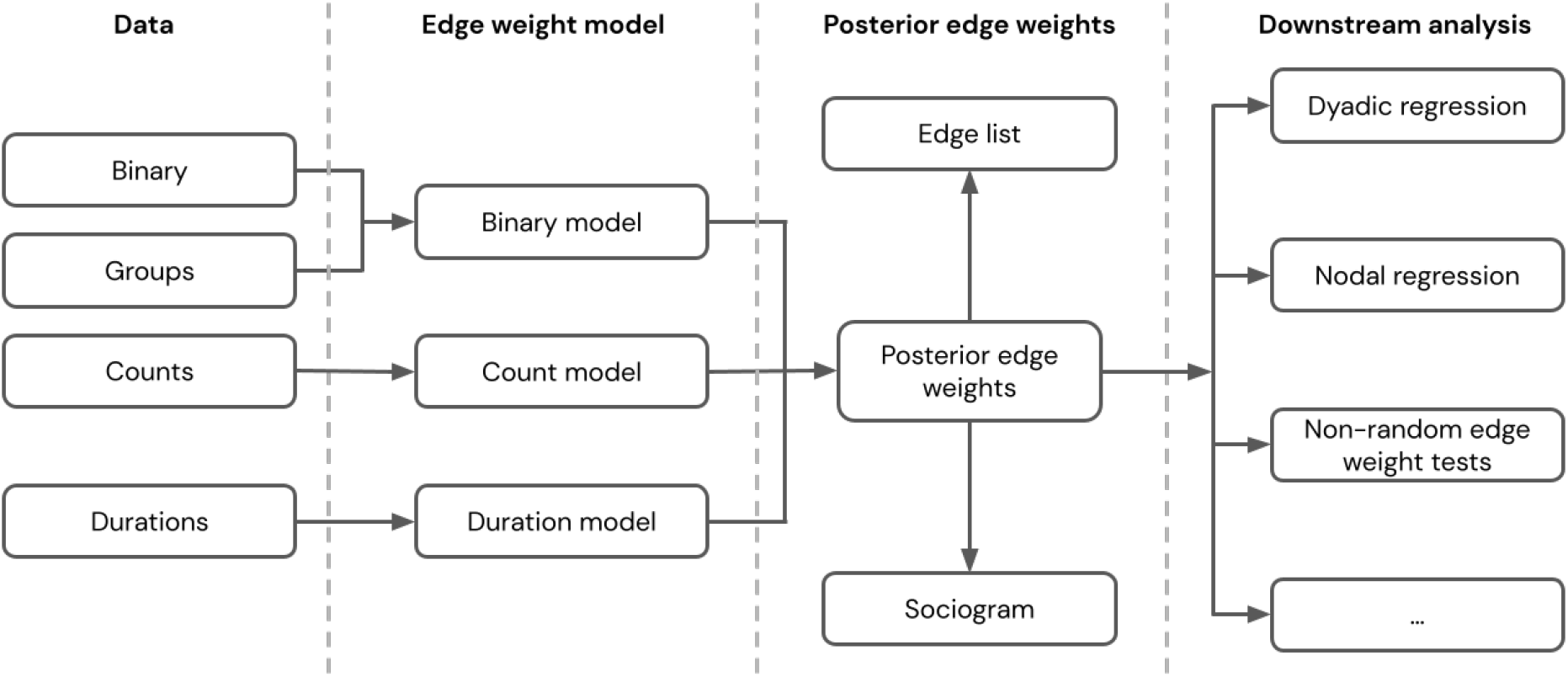
Schematic overview of the BISoN framework. Different types of observation data in various levels of aggregation are fed into the appropriate edge weight model for the type of data. The edge weight model generates posterior distributions over edge weights, quantifying uncertainty over them. Posterior edge weights can be visualised in a sociogram with uncertainty, or exported as an edge list with credible intervals. Posterior edge weights can also be exported for downstream analyses such as regression analyses, non-random edge weight tests, and more.

**Figure 2:**
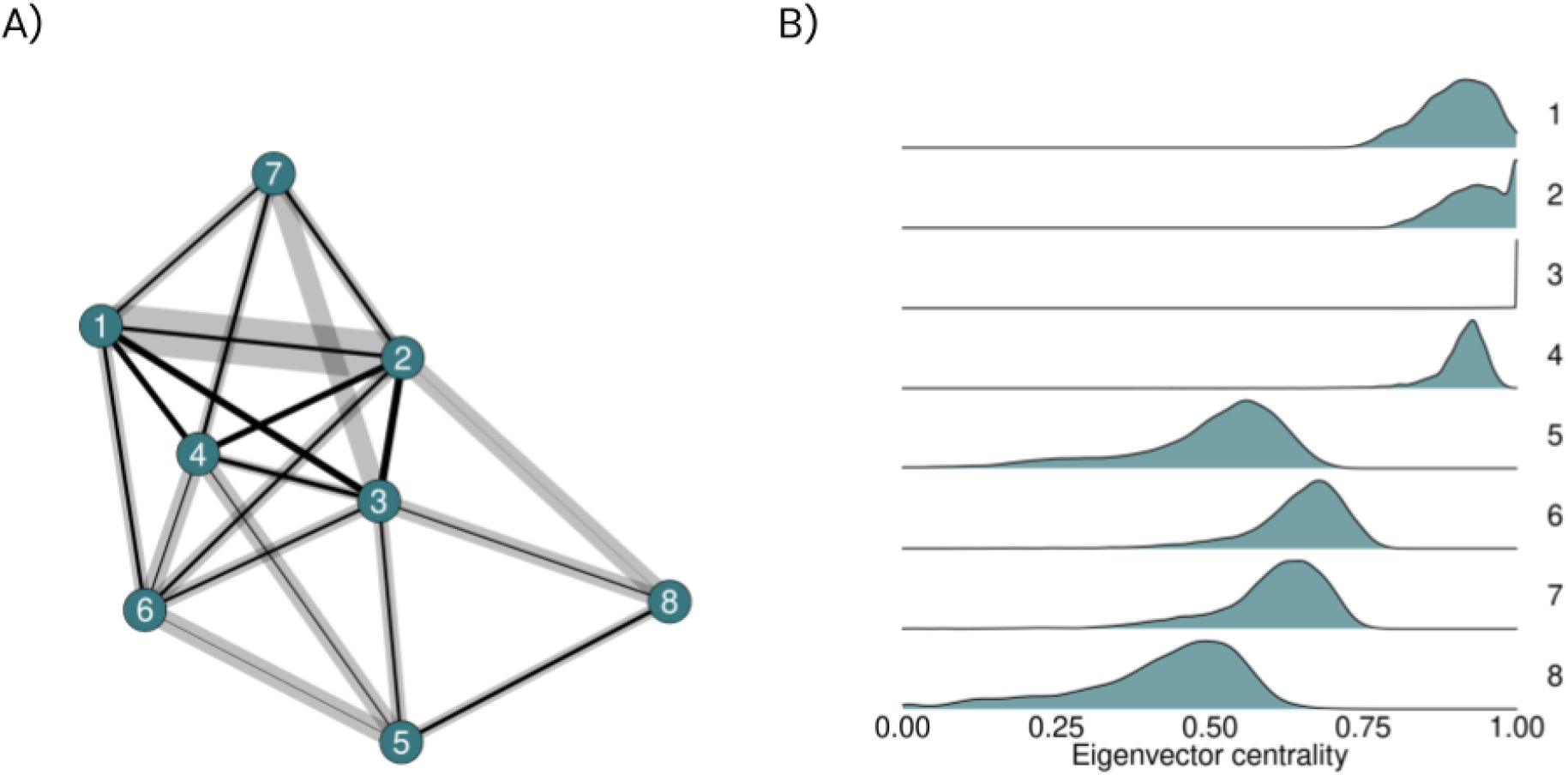
A) Visualisation of a network generated from uncertain edge weights. The width of edges denotes their weights, and the bands around the edges are proportional to the width of their 95% credible intervals. Low edge weights are arbitrarily removed from this plot (though not the analysis) for visualisation purposes. B) Posterior distributions of node-level eigenvector centralities for each node in the network. Nodes 1 - 4 are those assigned to the treatment, and nodes 5 - 8 to the control.

The posterior distributions of unstandardised eigenvector centralities for the 8 nodes are shown in Figure 2B. The treatment condition parameters are interpretable as the mean standardised eigenvector centrality of nodes for each of the two conditions. The 95% credible interval for the control parameter *β*_Control_ was [−1.27, −0.36], with a median of −0.83, and for the treatment parameter *β*_Treatment_ was [0.35,1.29], with a median of 0.85. One of the many benefits of Bayesian inference is that posterior distributions of different parameters can themselves be used gain statistical insights. In this case, the question of real interest is the difference between the two treatment conditions: *β*_Treatment_ – *β*_Control_. This quantity can simply be calculated by subtracting the posteriors corresponding to the treatment parameter from those corresponding to the control parameter. We found that the difference in standardised centrality between treatment conditions, *β*_Treatment_ – *β*_Control_, had a 95% credible interval of [0.99, 2.33] and a median of 1.69. The posterior distribution of the difference is also shown in Figure 2B. This indicates that the treatment condition appears to correlate strongly with a higher centrality.

This brief example outlines just some of the types of analysis that can be carried out in our framework. We have included several additional examples in the code included with this paper.

## Discussion

In this paper we presented BISoN, a general, highly flexible framework for modelling and analysing social networks. BISoN uses models of data collection processes to quantify uncertainty in edge weights. These models are versatile, and can be extended to capture additional social and non-social effects. We demonstrated how uncertainty in the network can then be propagated through to subsequent analyses, such as regressions involving network centrality. We have provided example code for how to implement the BISoN framework in R with Stan at: www.github.com/JHart96/BISON_examples.

BISoN acknowledges the inherent uncertainty in inferred edge weights, and by propagating this through downstream analyses can offer nuanced, balanced, and powerful statistical analyses, focused on quantifying biological effects and the uncertainty around them. As well as propagating uncertainty into downstream analyses, the edge weight model itself can also be adapted to ask questions about individual variation in gregariousness, reciprocation of directed behaviours, and underlying differentiation of edge weights, among many others (see the supplementary material for further information).

One of the main benefits of our framework is its inherent flexibility, but this also introduces a number of considerations when using it. Setting priors is an important part of Bayesian modelling, and it is important that they reflect realistic prior expectations of parameter values (van de Schoot et al., 2021). The priors included in the example code are not intended as good defaults, and instead the priors should be determined by careful consideration of the meaning of the parameter, prior expectations of likely values for the parameter, and using prior predictive checks (Conn et al., 2018). It is also important to note that using additional terms and altering the predictor part of the model can change the interpretation of model parameters, which may require the priors to be altered as well.

Another consideration is that network centrality measures based on binarised edges, such as degree and transitivity, have a slightly different interpretation in this framework. This is because, even though a particular dyad may never have been seen interacting in, say 100 samples, it is still possible that they would interact in the the 101st sample. The edge weight models BISoN uses naturally acknowledge this, so there is uncertainty around all edge weights. If binarised centralities are needed, there are two possible ways to use them in BISoN: 1) the edge weights can be thresholded, to generate a binary distribution on 0 and 1 for each edge; or 2) the edge weights can be assigned to categories, and membership of a specific category (e.g. weak or strong) becomes the new binary measure, again maintaining uncertainty over the membership. Option 1 can simply be applied to the sampling posterior by choosing a biologically relevant cut-off point and applying the threshold. In some cases this approach may prove challenging because it requires mapping domain knowledge and the question, in context, to a single threshold point for the analysis. Option 2 requires partial pooling over edge weights, assuming that they are drawn from a mixture of distributions, and assigns each edge weight to each type of connection with a certain probability. This approach is conceptually similar to the social bond categorisation method introduced by Ellis et al. (2021). If it is biologically sensible to believe there are different categories of connections in a system, this can provide a natural and principled way to compute binary network measures (see supplementary material for an example of how to do this).

A final consideration of our modelling framework is that, though non-social effects can be modelled in the predictor, this alone cannot remove some non-social effects, and could instead absorb the social effect of interest, depending on the underlying causal system. We recommend that these effects be treated with caution, and used only with explicit assumptions about the underlying causal system. These considerations are also discussed in further depth in the supplementary material using causal models.

The versatility of our proposed framework makes the approach powerful, but comes with a number of limitations. Though highly flexible, one important constraint on BISoN is that knowledge of sampling effort will be required to properly model the processes that generate observations. On a practical level, BISoN models will usually need to be fit using an MCMC sampler. MCMC can be slow and computationally expensive, especially with large datasets. If additional effects are not included, this issue can be partially overcome by aggregating data at the dyad-level and fitting collapsed versions of the models (see supplementary materials). Furthermore, MCMC methods do not guarantee convergence, so additional checks will be required to make sure the sampler has performed correctly (Cowles and Carlin, 1996; Draper, 2008). Together these limitations make the process of model building, fitting, and checking a more complex undertaking than simply building networks from point estimates. However, in return for these investments, researchers will have access to a powerful suite of tools that can make maximum use of hard-won data.

The BISoN framework is open-ended and can be extended in many ways. One particularly useful aspect of the Bayesian approach is the ability to treat data with uncertainty. This allows missing data to be modelled within the statistical model without the need for repeated model fitting routines. Furthermore, missing data can be estimated from other variables, which could be especially useful in populations where traits such as age are often estimated or unknown (Little and Rubin, 2002). Another potential direction for the framework is to move beyond static sociality measures and instead explicitly model the dynamics of edge weights, as discussed by Pinter-Wollman et al. (2014). Dynamic edge weights could be modelled by incorporating autoregressive time series models into the framework (Wei, 2013). This would make it possible to model the general trends in edge weights, how changes in edge weights propagate through the network, and even model flow processes such as disease or information transmission.

## Conclusion

Uncertainty affects all social network analyses, yet until now has been difficult to quantify. The BISoN framework we have introduced is designed to help quantify uncertainty and ensure it is preserved through subsequent analyses to give robust and reliable statistical inference. Our framework is general and widely applicable to almost any type of social relational data.

Estimated edge weights from our models can be used with many different types of downstream analysis while propagating uncertainty. We believe the ideas presented in this paper are only the first step in developing powerful, flexible, and robust statistical models for social networks. BISoN has the potential to enable us to ask a wide range of nuanced questions, and obtain a deeper insight into the nature of sociality.

## Author contributions

JDAH conceived of the initial idea behind BISoN, which was further developed with input from MNW, DWF, and LJNB. The example code was written by JDAH. The manuscript was written by JDAH with input from MNW, DWF, and LJNB.

## Acknowledgements

The authors would like to thank Rebecca Padget, Josefine Bohr Brask, Delphine De Moor, Samuel Ellis, Kenneth Keuk, Mario Angst, and members of the CRAB social network club for useful discussions on early versions of the manuscript. JDAH acknowledges funding from the Engineering and Physical Sciences Research Council [grant number EP/R513210/1]. LJNB acknowledges funding from a European Research Council Consolidator Grant (FriendOrigins - 864461) and from the National Institute of Health (R01AG060931, R01MH118203). DWF and MNW acknowledge funding from the Natural Environment Research Council [grant number NE/S010327/1]. DWF also acknowledges funding from the Natural Environment Research Council [grant number NE/S009914/1]. The authors declare no conflict of interest.

## Supplementary Material: Derivation of duration model

With duration data, edge weights are often calculated as:

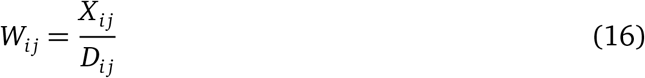

where *D_ij_* is the maximum amount of time *i* and *j* could have been seen engaged in a social event. The total amount of time *X_ij_* is the sum of times from *K_ij_* social events:

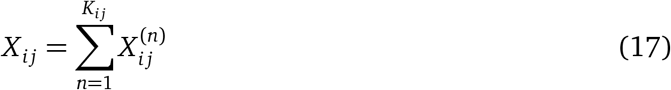

where 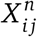 is the *n*-th observed social event between *i* and *j*, and *K_ij_* is the number of social events between *i* and *j* over the course of the study period.

This means that edge weights could alternatively be modelled in terms of the mean rate of events per unit time, *λ_ij_* and the mean event duration *μ_ij_*, as follows:

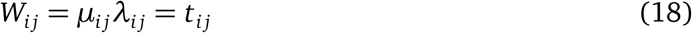

where *t_ij_* is the proportion of time *i* and *j* spend engaged in a social event.

Unlike the binary and count models, there is no natural process model for duration data. This leaves the modelling decisions for *λ_ij_* and *μ_ij_* less clear. Therefore we introduce a minimal process model that makes few assumptions but still attempts to capture as much detail about the generating process for the data as possible. Time-until-event processes are often modelled as exponential processes. We propose to use the exponential distribution, centered on the mean event duration *μ_ij_*, to model the duration of events:

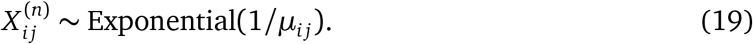

Additionally, the count of events can naturally be modelled as a Poisson process, with rate *λ_ij_*. Combining these assumptions gives the following model:

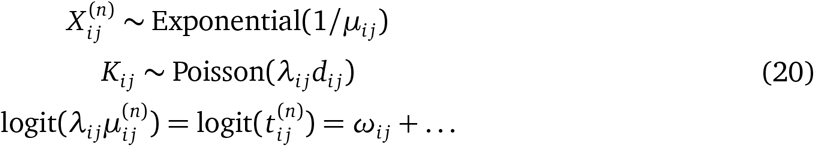

Unfortunately this model is challenging to fit because it requires introducing a latent variable to describe 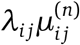. This latent variable can be marginalised out by using the equality 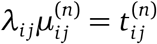, giving an alternative parameterisation of the model:

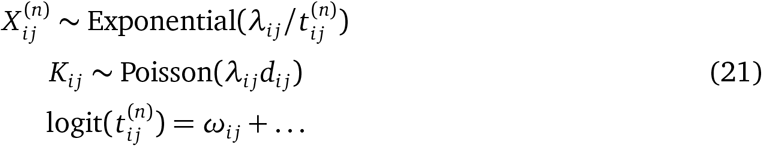

In this specification, it is theoretically possible for the duration time 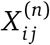 to exceed the maximum possible duration time. A truncated exponential distribution could be used to prevent this from occurring, but for simplicity we have presented a simpler model here. There are many alternative possible specifications for the duration model, so predictive checks will be especially important when using these kinds of model.

## Supplementary Material: Non-random edge weight test for duration data

The non-random edge weight test compares two alternative edge weight models, one where social events between dyads depend on a dyad-level edge weight, and one where they depend on a single, population-level edge weight. This tests the hypothesis that apparent differences in edge weight between dyads is due only to sampling, or if there there is an underlying difference in edge weights. For duration data, the two models are described by:

**H1**

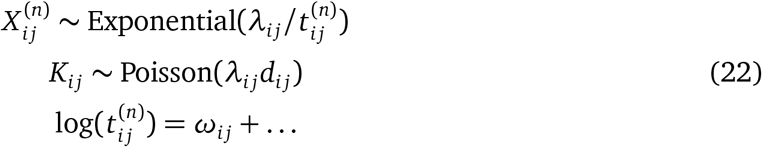

**H0**

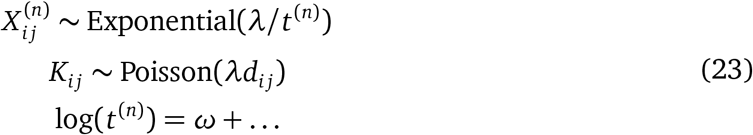

## Supplementary Material: Modelling individual gregariousness

The base edge weight model uses a single term to model edge weight, *ω_ij_*, where observed social events depend on it in some way, for example in the binary model:

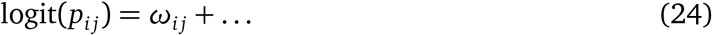

where … represent additional control terms for nuisance effects/confounds. This is designed to capture a dyad-level measure of edge weight, but depending on the question and context, it may be necessary to consider additional effects of edge weight, for example node-level gregariousness. This could be modelled quite simply by including a node-level term in the above equation:

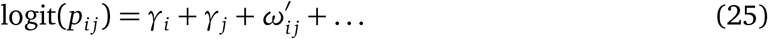

where *γ_i_* is a latent measure of node-level gregariousness, and 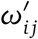 is a measure of the edge weight of a dyad beyond that explained by individual gregariousness. The original edge weight from the edge weight model can be recovered as 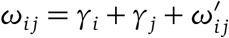.

This alternative formulation may be useful if individual gregariousness is of particular interest.

## Supplementary Material: Including an intercept term in the predictor

In some cases it may be useful to include an ‘intercept’ term in the predictor part of the edge weight model. For example, in the simple version of the edge weight model for binary data, the predictor:

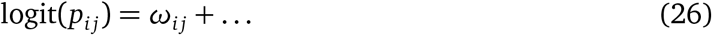

could be modified to:

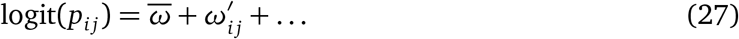

where 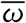 is the mean log-odds edge weight of all the dyads, and 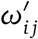 is a zero-centered relative measure of edge weight. This measure is equivalent to taking the standard edge weight 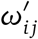 and mean-centering it.

When including an intercept term, the intercept parameter will capture the mean edge weight, leaving the *ω_ij_* term to explain differences between the *ij*-th edge and the mean. Using this formulation, the edge weights might then be interpreted relatively as *positive* or *negative*, relative to the mean. This could lend novel insights into the structure of networks.

## Supplementary Material: Sharing information between dyads with partial pooling

In some cases it can be useful to assume that edge weights are drawn from the same distribution. This assumption allows information about edge weights between dyads to be shared, and decrease the uncertainty around estimates. This may not always be the case, but the assumption can be checked using predictive checks. This is an important consideration for dyads that are rarely seen in the same sampling period. There is little information for these dyads and they will be heavily influenced by any type of pooling. Depending on the accuracy of the pooling assumption, this could greatly improve the inference. Predictive checks should be used when determining the appropriate model for any given dataset.

Partial pooling of edge weights can be implemented by adding an adaptive prior to the edge weight terms. Using the count model as an example, a model using unimodal partial pooling would take the form:

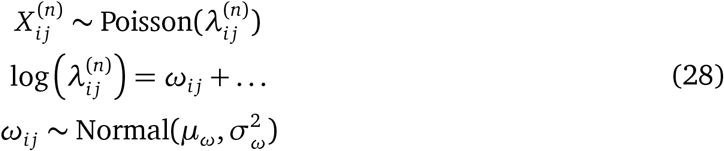

where *μ_ω_* and *σ_ω_* are the mean and standard deviation of the adaptive prior over edge weights.

Edge weights are not necessarily unimodally distributed, and may comprise multiple different types of edge weight. Mixture models have previously been used to study social complexity, but they can also be incorporated into our framework to capture the multimodal nature of edge weights. A model using multimodal partial pooling would take the form:

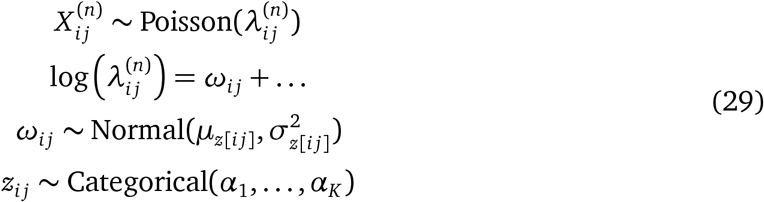

where *Z_ij_* is an index variable describing the mixture component to which each dyad belongs. The *α_k_* parameters describe the weightings of each mixture component, centered on the component with mean *μ_k_* and standard deviation *σ_k_*.

## Supplementary Material: Directed networks and reciprocal effects

The social relations model often used in human social network studies includes a provision for reciprocal effects using correlated effects for sending and receiving effects (Kenny and La Voie, 1984). These phenomena have also been modelled in more recent developments of the framework, such as in the STRAND framework (Ross et al., 2022). In undirected networks there is no need for modelling reciprocal effects, but it may be useful to do so in directed networks for two reasons: 1) reciprocal effects may be of interest for the question in hand, and 2) including a reciprocal effect may improve inference and reduce uncertainty around parameter estimates if there is in fact a reciprocal effect at play.

To include reciprocal effects in the base edge weight model, the prior for the directed edge weights can be replaced with a multivariate normal distribution:

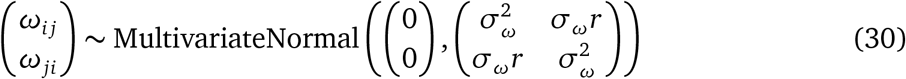

where *σ_ω_* is the standard deviation of edge weights, and *r* is the correlation between sending and receiving effects. The correlation parameter *r* can be interpreted as the correlation between sending effects and receiving effects. If the correlation is positive this would indicate that directed social events tend to be reciprocated, whereas a negative correlation implies that social events are imbalanced.

## Supplementary Material: Discussion on controlling for nonsocial effects

There has been considerable attention in the social network literature focusing on methods for controlling for non-social effects in statistical analyses. The key concern is that edge weights may be affected by non-social factors, for example preferences for certain spatial locations, rather than preferences for specific individuals. This is of obvious importance when conducting social network analyses, as it could cause the wrong conclusions to be drawn from statistical models.

A number of methods have been proposed to attempt to control for non-social effects, also termed nuisance effects or confounds. Recent works have focused on computing correction terms to remove the effect of non-social factors from analyses (Farine and Carter, 2021; ?). Our framework also introduces a mechanism for controlling for non-social effects. However, there are implicit limitations to all of these methods (including BISoN) because of the nature of the underlying causal systems. Here we present a few examples in a causal framework using directed acyclic graphs (DAGs) to illustrate the considerations when attempting to control for non-social effects. DAGs and causal inference are a powerful tool for reasoning over causal problems, for those unfamiliar with causal inference, Pearl et al. (2016) provides a thorough primer.

The DAG shown in Figure 3 describes the standard causal system used for inferring sociality in social network analysis. Consider the hypothesis that the trait *T* of a dyad (e.g. age difference) affects the dyad’s sociality, *S*. Unfortunately, *S* is unknown and unknowable, but we assume it is borne out in observable social events *X*, which can be measured in any number of ways. *X* is therefore a good measure of sociality, and can be used to ask questions about the relationship between *T* and *S*.

**Figure 3:**
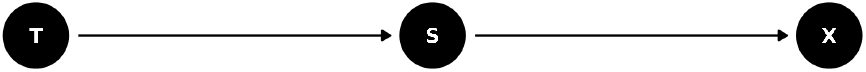
*T* influences *S*, which influences *X*. *X* is only affected by *S*, so it can serve us well as a proxy for sociality.

### Example 1

Problems can start to arise when there is an additional effect on *X*, a non-social effect *N*. The first example we will consider is shown in Figure 4. In this diagram, *T* influences *S*, which in turn influences *X*. *X* is also influenced by *N*, causing a non-social effect on *S*. *N* and *S* are independent, so using *X* as a measure of *S* will give an unconfounded, but imprecise measure of sociality. Alternatively, if *N* can be controlled for, *X* will be a more precise measure of sociality. In this example, controlling for the non-social effect is not essential, but will yield more precise estimates and improved inference.

**Figure 4:**
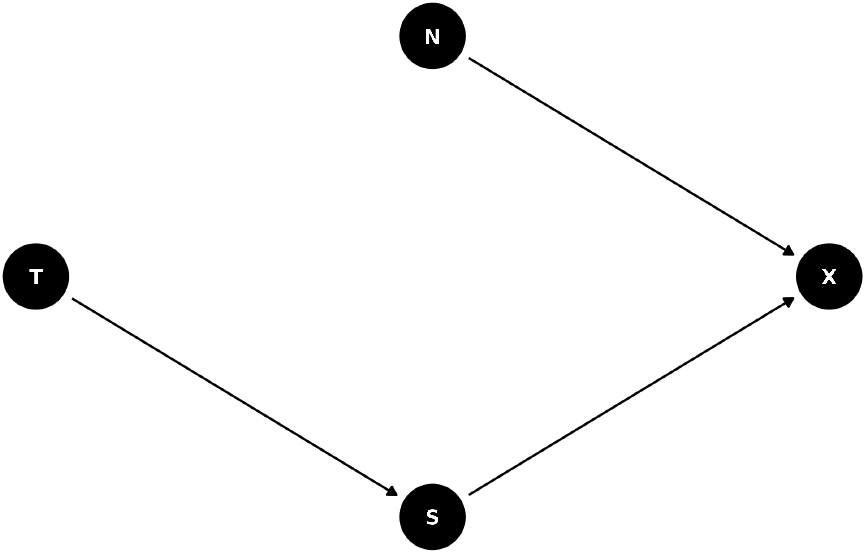
DAG 1: *T* influences *S*, which in turn influences *X*. *X* is also influenced by *N*, causing a non-social effect on *S*. *N* and *S* are independent, so controlling for the non-social effect is not essential, but will yield more precise estimates and improved inference.

### Example 2

The second example we will consider is shown in Figure 5. *T* influences *S*, which in turn influences *X*, but *T* also influences *N*, which in turn influences *X*. *N* and *S* are now independent, conditional on *T*. Using *X* as a proxy of *S* will no longer work. The relationship between *T* and *X* is now confounded by *N*, so *N* must be controlled for before *X* can be used as a measure of sociality. Controlling for *N* essentially blocks the causal influence of *T* on *X* through *N*, so *X* becomes a good measure of sociality only when *N* is controlled for.

**Figure 5:**
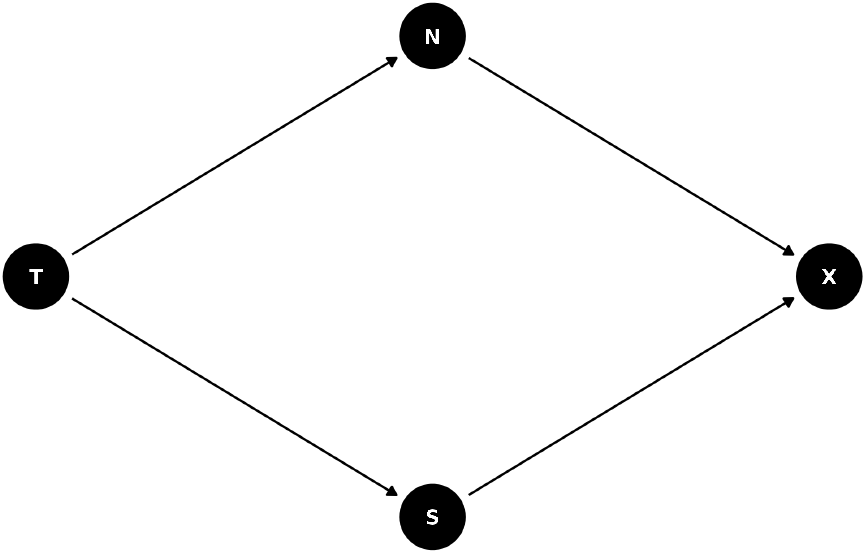
DAG 2: *T* influences *S*, which in turn influences *X*, but *T* also influences *N*, which in turn influences *X*. *N* and *S* are still independent, but now using *X* as a proxy of *S* will no longer work. The relationship between *T* and *X* is now confounded by *N*, so *N* must be controlled for before *X* can be used as a measure of *S*.

### Example 3

The final example we consider is shown in Figure 6. In this example *T* influences *S*, which in turn influences *X*, and again *T* also influences *N*, which in turn influences *X*. This time, however, *N* and *S* are no longer independent, conditionally or otherwise, there is now a direct influence of *S* on *N*. This means that *T* influences *X* on two distinct paths: *T* → *S* → *X*, and importantly, *T* → *S* → *N* → *X*. The overall influence of *T* on *X* can no longer be parsed out from the effect of *N* on *X*. There is no way to separate out the social and non-social effects without additional information.

**Figure 6:**
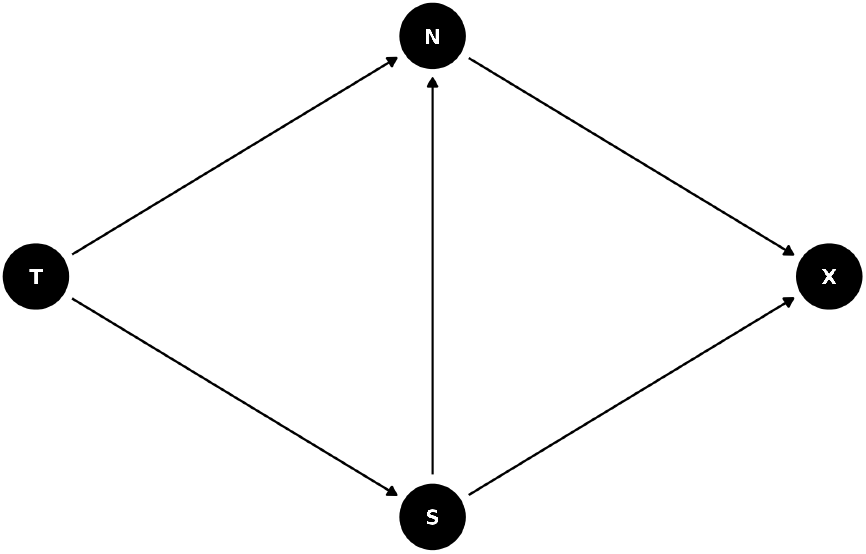
DAG 3: *T* influences *S*, which in turn influences *X*, and again *T* also influences *N*, which in turn influences *X*. There is a direct influence of *S* on *N*, preventing social and non-social effects on *X* from being separated.

This example might arise where *X* is derived from a spatial measure, *N* is a spatial variable, and *T* affects both sociality and spatial preference. Since sociality requires individuals to spent time in the same location, but the trait also affects location preference, the non-social spatial preference cannot be parsed out from the edge weight. Including the non-social effect as a control here would remove one of the paths for *T* to influence *X*, and could remove the effect of interest, but excluding the non-social effect as a control opens up the possibility for a nonsocial effect to be mistaken as a social effect.

### Modelling non-social effects

These examples show the potential pitfalls when controlling for non-social effects. In examples 1 and 2, controlling for the non-social effect is always useful and improves the precision of estimates while removing the effect of unimportant non-social effects. However in example 3, including the control could remove the effect of interest when it is there, but excluding the control could deceive us and detect an effect of interest when there is none.

## Supplementary Material: Models for aggregated data (collapsed models)

### Aggregated binary model

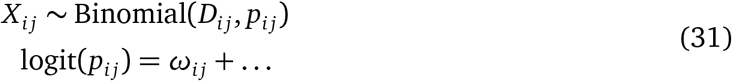

### Aggregated count model

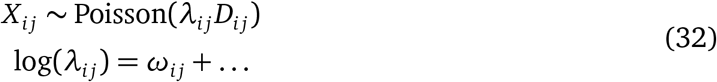

### Aggregated duration model

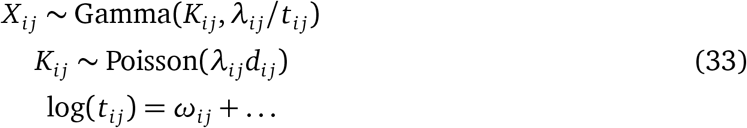

## Supplementary Material: Modelling group-based data with the binary model

A popular method of data collection in fission-fusion societies is “gambit of the group”. This method of data collection assumes individuals gather in groups according to their social network, so a sighting of three individuals A, B, and C in a group or gathering counts as a social event between A and B, between A and C, and between B and C. Since the observations are either presence or absence of a social event, the binary model is an appropriate model for these data. However, since social events derived from the same grouping are unlikely to be independent, we propose to include a varying effect to control for group membership, as follows:

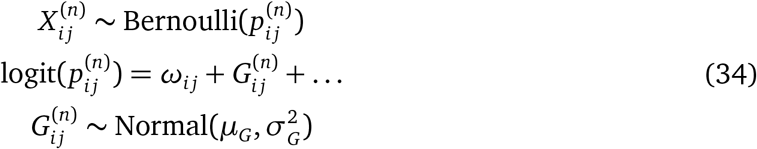

where 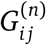 is the group-level effect for the *n*-th grouping that *i j* were seen in. This parameter will be shared by any observations for dyads in the same grouping. An example of this type of model is included in the example code. Additionally, the convert_gbi.Rmd file gives an example of how group-by-individual matrices can be converted into the long format required by Stan.

The long format used to fit these types of models can generate large numbers of observations depending on the number of individuals and groupings. Be aware it may be computationally intensive to process the data and fit the model. We have included an example of model fitting using INLA.

## Supplementary Material: Fast Inference with INLA

Some configurations of BISoN models belong to a class of models that can be estimated well using integrated nested Laplace approximation, a computational technique that can approximate posterior distributions of parameters as Gaussian distributions (Rue et al., 2017). The R package R-INLA can be used to fit popular types of model such as GLMMs, so to use INLA with BISoN, the BISoN models need to be reformulated in terms of GLMMs (Rue et al., 2009). See the Github repository for full examples on how to use INLA with BISoN (www.github.com/JHart96/BISON_examples).

### Binary data edge weight model

The edge weight model for binary data described above can be extended in several ways, but one of the main general forms is the following:

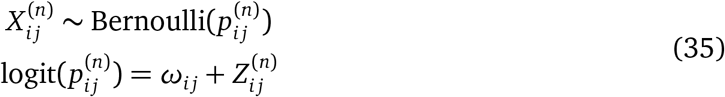

where 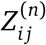 is a varying effect (also known as a random effect) for some aspect of the observation of the *i j*-th dyad at the *n*-th observation. This form of the model is equivalent to a generalised linear mixed model with the binomial family. The model code for fitting this model is:

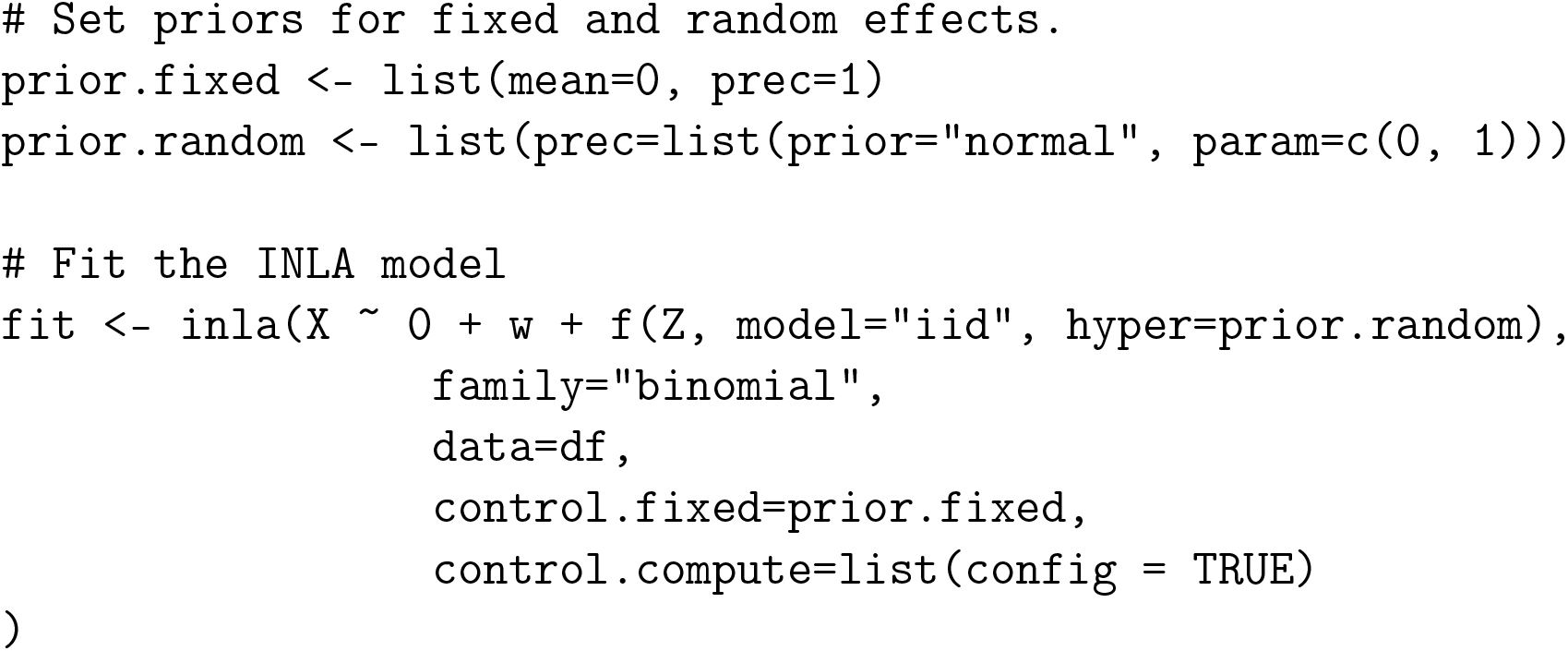

### Count data edge weight model

The simple configuration of the edge weight model for count data is not exactly equivalent to a GLMM, as the rate parameter 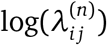 is multiplied with the observation time 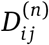, shown in this example below:

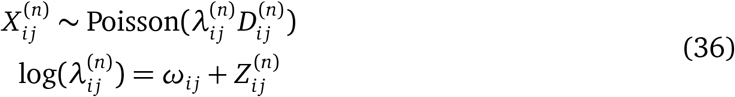

The standard Poisson family GLMM models the mean count, not the mean rate, so an *offset* must be used to correct for observation time 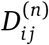 The model can be fitted in R-INLA using:

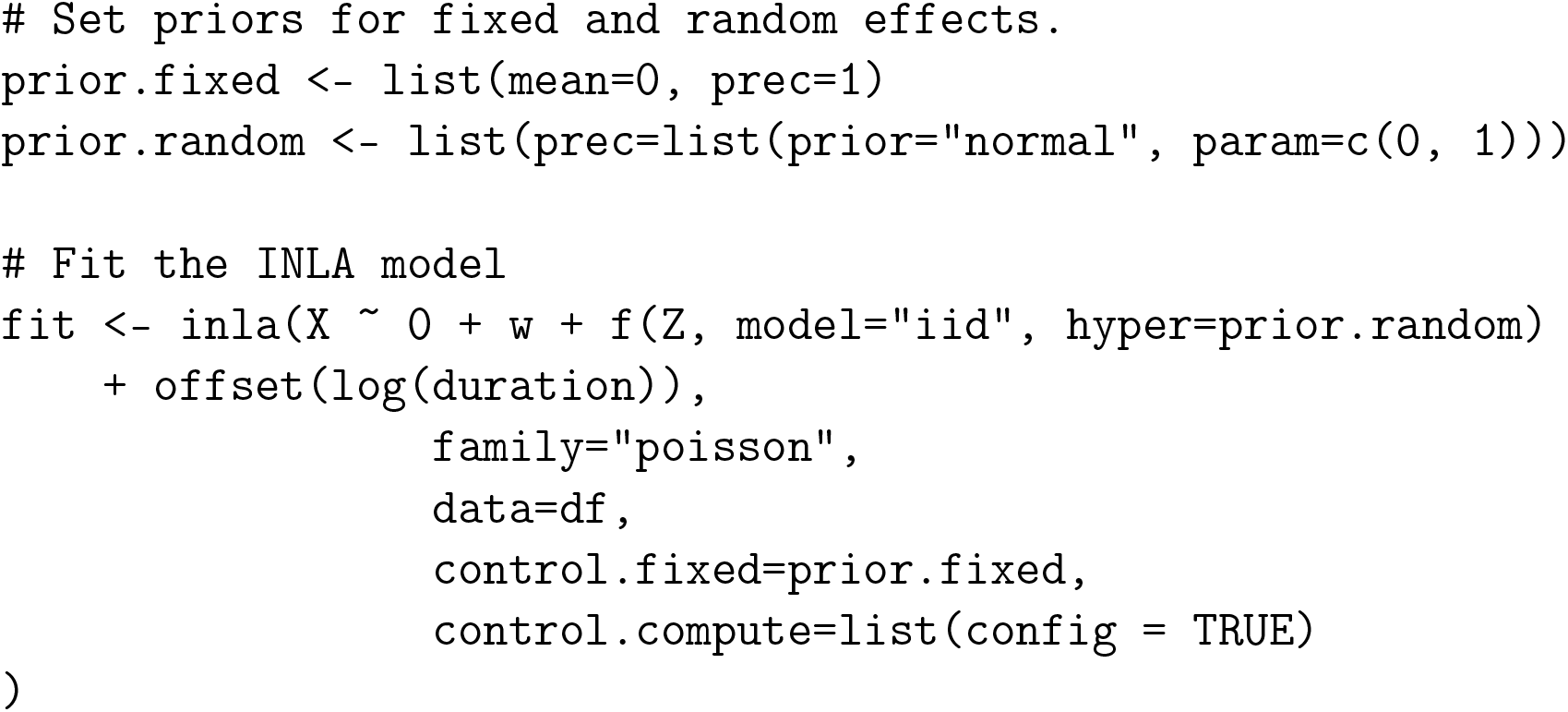

### Post-processing

When using INLA, the edge weight posteriors *ω_ij_* are estimated as univariate normal distributions, and be extracted from a fitted model using the following code:

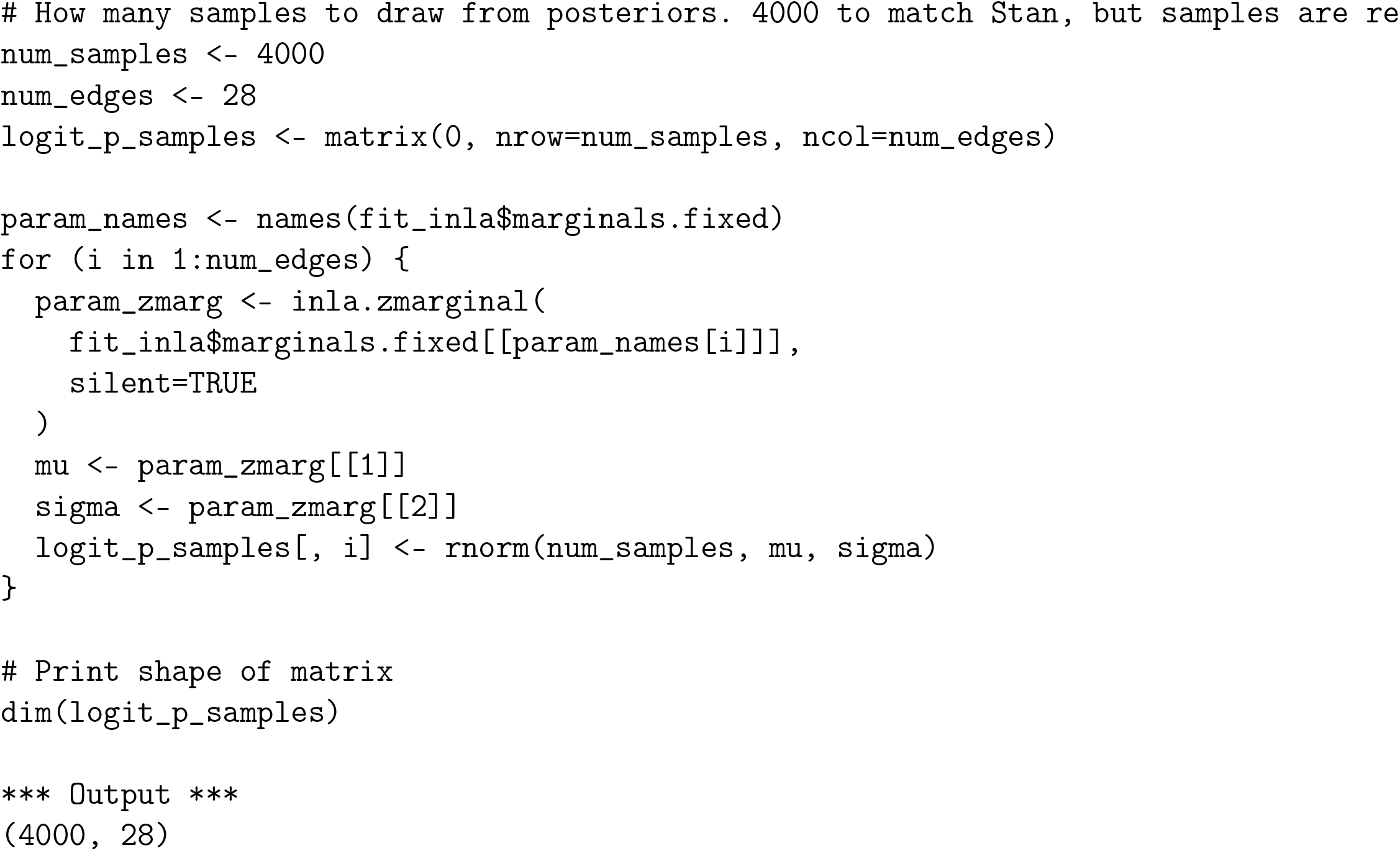

